# Effects of arousal and movement on secondary somatosensory and visual thalamus

**DOI:** 10.1101/2020.03.04.977348

**Authors:** Gordon H. Petty, Amanda K. Kinnischtzke, Y. Kate Hong, Randy M. Bruno

## Abstract

All neocortical sensory areas have an associated primary and secondary thalamic nucleus. While the primary nuclei encode sensory information for transmission to cortex, the nature of information encoded in secondary nuclei is poorly understood. We recorded juxtasomally from neurons in secondary somatosensory (POm) and visual (LP) thalamic nuclei of awake head-fixed mice with simultaneous whisker tracking and pupilometry. POm activity correlated with whether or not a mouse was whisking, but not precise whisking kinematics. This coarse movement modulation persisted after unilateral paralysis of the whisker pad and thus was not due to sensory reafference. POm continued to track whisking even during optogenetic silencing of primary somatosensory and motor cortex and after lesion of superior colliculus, indicating that motor efference copy cannot explain the correlation between movement and POm activity. Whisking and pupil dilation were strongly correlated, raising the possibility that POm may track arousal rather than movement. LP, being part of the visual system, is not expected to encode whisker movement. We discovered, however, that LP and POm track whisking equally well, suggesting a global effect of arousal on both nuclei. We conclude that secondary thalamus is a monitor of behavioral state, rather than movement, and may exist to alter cortical activity accordingly.

## Main

Somatosensory, visual, auditory, and gustatory cortex are each reciprocally connected with a specific subset of thalamic nuclei. These nuclei can be subdivided into primary and secondary (often termed “higher-order”) nuclei^1–3^. The primary nuclei are the main source of sensory input to the cortex and respond robustly to sensory stimulation with low latency^4–7^. Unlike primary nuclei, the secondary nuclei are interconnected with many cortical and subcortical regions, and their role in sensation and cognition is poorly understood.

In rodents, the facial whisker representation of primary somatosensory cortex (S1) is tightly integrated with two thalamic nuclei: the ventral posteromedial nucleus (VPM) and the posterior medial nucleus (POm). Compared to the primary nucleus VPM, the secondary nucleus POm has broader receptive fields, longer-latency sensory responses, and poorly encodes fine aspects of whisker touch such as contact timing and stimulus frequency^8–11^. It receives input from S1, motor cortex, posterior parietal cortex, the zona incerta, and many other subcortical regions in addition to brainstem afferents^4,12,13^. Whereas VPM innervates cortical layer 4 and the border of layers 5B and 6, POm projects to the apical dendrites of layer 1 as well as layer 5A^6^. POm is a stronger driver of layer 2/3 cells than cortico-cortical synapses and can enhance sensory responses in pyramidal neurons of layers 2/3 and 5^14,15^. POm is thus positioned to strongly influence sensory computations in S1 and do so in ways that are highly distinct from VPM. However, what POm activity encodes remains a mystery.

One possibility is that POm activity encodes self-generated movements, through either sensory reafference (stimulation of the sensory receptors by active movement) or motor efference copy (internal copies of motor commands), rather than extrinsic tactile sensations^16^. If secondary thalamus were a monitor of movements^5^, somatosensory cortex could use POm input to differentiate self-generated and externally generated sensory signals. However, recent studies in awake animals have observed that, in comparison to VPM, POm poorly encodes whisker motion and contact^8,17^, which casts doubt on the hypothesis that secondary pathways provide detailed motor information to cortex.

An alternative hypothesis is that secondary thalamus is a key structure for monitoring behavioral state. For instance, several studies have noted that a subset of POm neurons are activated by pain^18,19^, a powerful stimulus that can trigger a change in animal’s state. Spatial attention is a more subtle form of behavioral state change and has been implicated repeatedly in studies of primate secondary visual thalamus (lateral pulvinar)^20–22^. The rodent homolog to the pulvinar (lateral posterior nucleus, LP) is active during mismatch of movement and visual stimuli^23^, which might reflect elevations in visual attention or even global arousal. These results raise the possibility that modulation by behavioral state is a general feature of all secondary nuclei.

Here we investigate how afferent, corticothalamic, and collicular inputs– the three main excitatory pathways to secondary sensory thalamus—influence encoding of movements by POm in the awake mouse. We discovered that removing these circuits enhances rather than reduces modulation of POm activity by movements, suggesting that these pathways may mainly transmit signals of a nature other than movement. We further examine how POm activity compares to that of LP – which have not been directly compared before - to investigate general principles of secondary thalamus function. This comparison reveals that behavioral state, rather than movement itself, prominently dictates the activity of secondary thalamus.

## Results

We characterized the degree to which POm encodes whether or not an animal is whisking versus the fine details of whisking movements. We recorded juxtasomally from single neurons in head-fixed mice while acquiring high-speed video of the contralateral whisker field, from which whisker positions could be algorithmically extracted (Figure 1a, b)^24^. To measure slow aspects of whisking, we calculated whisking amplitude from the median angle of all whiskers. Whisking amplitude is defined as the difference in angle between minimum and maximum protraction over the whisking cycle (Hill *et al.*, 2011; Moore *et al.*, 2015, see Methods). Whisking amplitude was then used to determine periods of quiescence and whisking, as defined by periods of time when whisking amplitude exceeded 20% of the maximum for more than 250 msec (Figure 1b, shaded areas).

**Figure 1.**
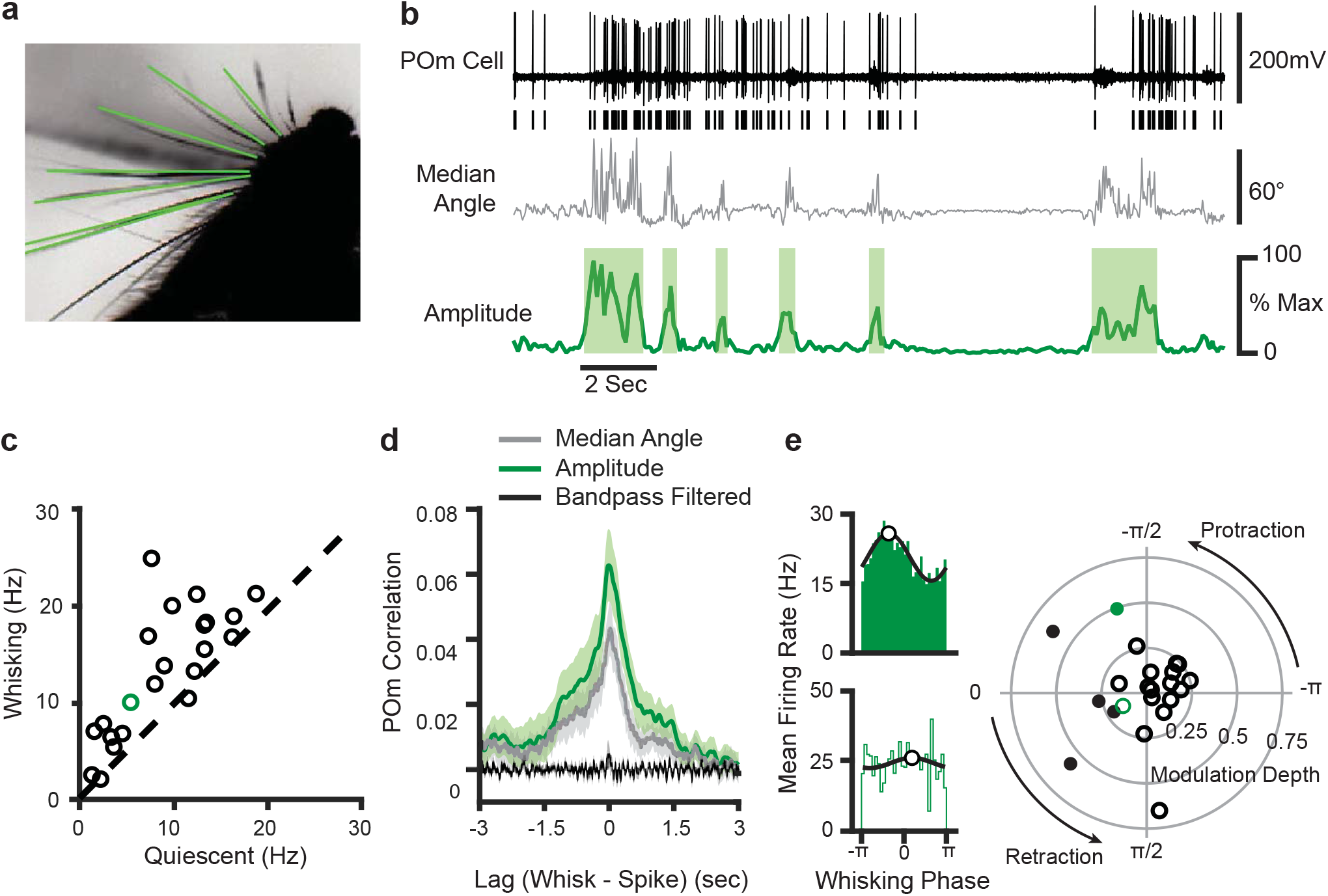
POm cells mainly track slow components of whisking activity. **(a)** An example frame from a video, captured at 125 FPS. Identified whiskers are highlighted in green, and whisker bases are indicated by yellow circles. **(b)** Example traces of juxtasomal POm recording and whisking. The median angle of all whiskers in each video frame (middle, gray) was used to calculate the whisking amplitude (bottom, green). **(c)** Scatter plot of POm firing rates during whisking and quiescence (n = 22 POm cells, 5 mice, increase from mean of 7.8Hz to 12.4 Hz, or 58%, p < 10^−4^, paired t-test). *Green*, cell in 1b. **(d)** Cross-correlation of POm firing rate and whisking amplitude (green), angle (gray), and 4−30 Hz bandpass-filtered angle (black). Shading, SEM over cells. Cross-correlation is normalized such that autocorrelations at zero lag equal one. **(e)***Left,* Firing rate as a function of phase in the whisking cycle for two example POm units. A sinusoid model (black) was fit to each cell to quantify preferred phase (white markers) and modulation depth. *Right,* A polar plot of modulation depth (radius) and preferred phase (angle) of each POm unit. *Filled circles*, cells with significant phase modulation (p<0.05, Kuiper test, Bonferroni corrected).

Whisking substantially elevated POm firing rates. We computed the mean firing rate for each cell during periods when the mouse was whisking versus quiescent (Figure 1c, 22 POm neurons in 5 mice). The firing rates of POm cells were significantly higher during bouts of whisking, increasing from a mean firing rate of 7.8Hz to 12.4 Hz (58.5% increase, p < 10^−4^, paired t-test). To understand which components of whisking might drive POm activity, we calculated the cross-correlation between POm firing rate and three features of whisking activity: the median angle (Figure 1d, gray), the amplitude metric which captures the slow envelope of whisking (green), and the median angle bandpass-filtered from 4−30 Hz (black), which reflects fast protractions and retractions of the whisker. We found that POm neurons had little correlation with the bandpass-filtered angle, but prominent correlations with both whisker angle and whisking amplitude around a time lag of zero. The strongest correlate of POm activity was whisking amplitude, suggesting that POm is coupled to the slow components of whisking, rather than tracking individual whisk cycles.

To further investigate the encoding of the fast components of whisking in POm, we analyzed whether individual cells preferred to discharge during a certain phase of the whisking bout. We quantified the phase of whisking by applying the Hilbert transform to the bandpass-filtered median whisker. We identified the phase at which each action potential occurred during whisking and plotted distributions of firing rate as a function of phase. For each cell, we fit a sinusoid to characterize the cell’s preferred phases (the phase of the whisk cycle that elicited the highest firing rate) and modulation depth (the degree to which phase impacts firing rate, measured as the peak-trough difference normalized by mean firing rate). Figure 1e shows the phase relationship of two example cells: one with significant coding (top) and the other insignificant (bottom). Most POm cells (17/22) resembled the non-phase coding example, having little or no modulation (right). Together, these results indicate that the majority of POm cells do not encode fast whisking dynamics such as whisker angle or the phase of the whisking cycle. Rather, they track overall whisking activity, *i.e.* the whisking versus quiescence.

One possible source of whisking-related activity is reafferent sensory input: when the mouse whisks, the self-generated movement could deform the whisker follicle and stimulate mechanoreceptors. To measure the degree to which POm activity is driven by the sensory reafference caused by whisking, we severed the facial motor nerve on the right side of the face (Figure 2a), contralateral to our recordings, while taking video of the left (ipsilateral) side of the face. This manipulation does not damage the sensory neurons and avoids the risk of inducing sensory neuron plasticity (Shetty, Simons, 2003). Mice were no longer able to move the right whisker pad, but the whisking on the left side of the face was unaffected (Figure 2b). Without whisker movement, there can be no reafferent sensory input from the right whisker pad. As in intact mice, firing rates of POm neurons in nerve cut animals were significantly higher during whisking bouts (Figure 2c, n = 12, p = 0.0007, paired t test) and to a similar degree, increasing from quiescent firing rate of 11.6Hz to a whisking rate of 16.7Hz (44%). POm firing rates also correlated with ipsilateral whisking amplitude, at a similar magnitude and with a similar lag as in the contralateral whisker field in intact mice (Figure 2d). This demonstrates that the correlation of POm activity and overall whisking is not due to ascending reafferent information.

**Figure 2.**
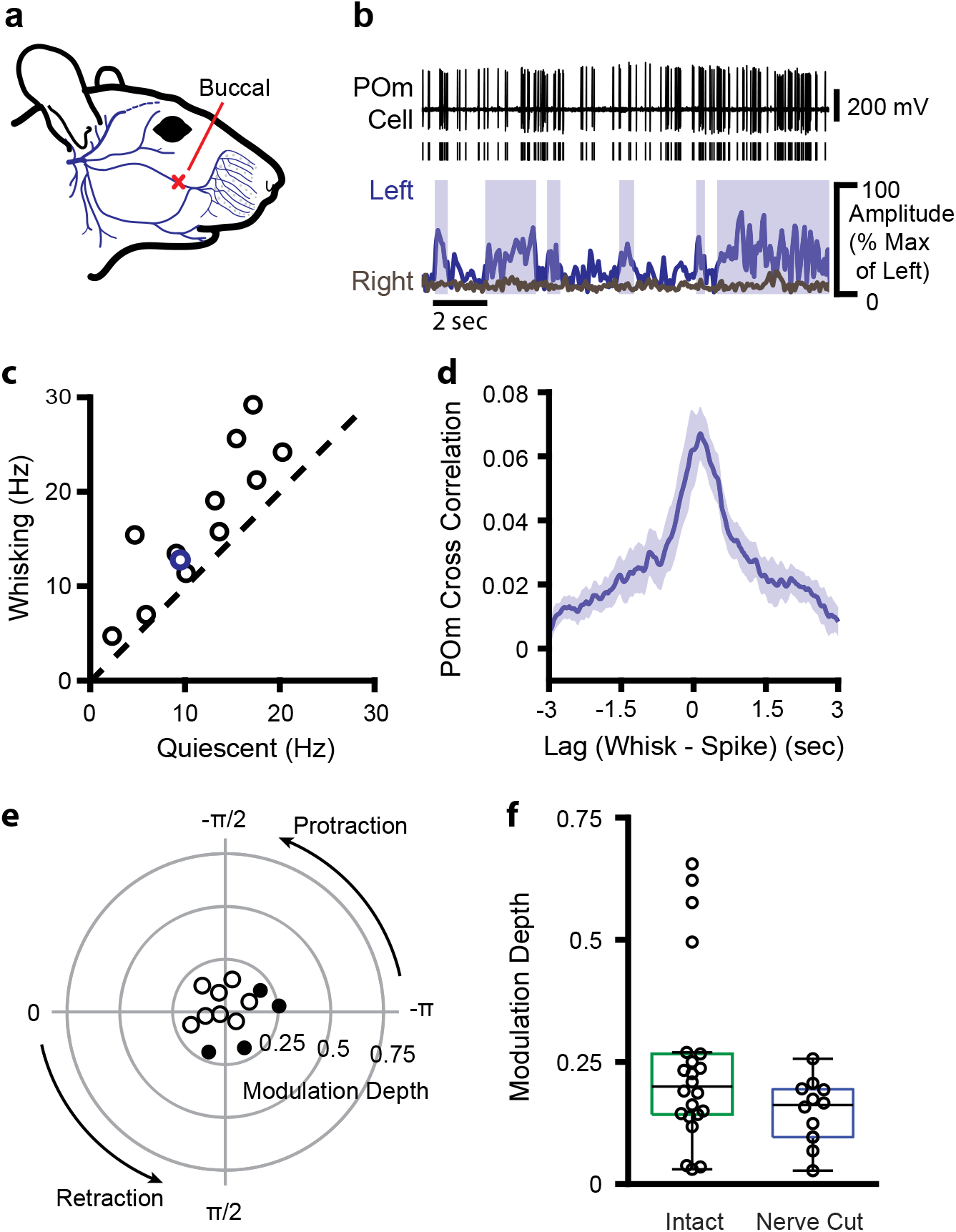
POm encodes whisking activity in absence of reafferent sensory input. **(a)** The buccal branch of the facial motor nerve was severed unilaterally, preventing whisker motion on the right side of the face. Adapted from Heaton *et al.*, 2014^56^. **(b)** Example POm cell (top, black), ipsilateral (left side of face) whiskers (bottom, blue), and contralateral whiskers (*bottom, gray*). Blue boxes: periods of whisking as in Fig.1B. **(c)** Scatter plot of mean POm firing rate during whisking and quiescence. *Blue*, example cell in B. Firing rates during whisking are significantly higher than quiescence (n = 12 cells from 2 animals, quiescent mean: 11.6Hz, whisking mean: 16.7Hz, 44% change, p = 0.0007, paired t test). **(d)** Cross-correlation of POm firing rate and ipsilateral whisking amplitude. **(e)** Polar plot of modulation depth and preferred phase of each POm unit as in Figure 1E. **(f)** Modulation depth of POm cells in intact mice (green, as in Figure 1E) and after buccal nerve cut (blue). There was a significant difference in the variance of modulation depth between groups (p = 0.0013, two-sample F test).

We also calculated the phase coding of the ipsilateral whisker field (Figure 2e) and compared it to phase coding of the contralateral whisker field (Figure 1e). While average modulation depth was unchanged (Figure 2e p = 0.12, Wilcoxon rank-sum test test), modulation depth by definition is bounded at zero, complicating analysis of medians close to zero. Indeed, there was a noticeable and statistically significant decrease in the range of modulation depths in the transected group (Figure 2f, p = 0.0013, two-sample F-test), consistent with nerve cut eliminating the largest modulation values. These results suggest that reafferent signals do not contribute to changes in POm activity reflecting the slow envelope of whisking (Figure 2d) but are responsible for the small population of POm cells carrying fast phase information (Figure 2e).

POm is reciprocally connected to several cortical areas, potentially making cortical input a strong driver of POm activity^4,15,26^. In particular, input from S1 and primary motor cortex (M1) could convey sensorimotor information, such as a motor efference copy, that would drive whisking-related activity, independent of ascending sensory input. To test this, we expressed halorhodopsin in all excitatory cortical neurons by crossing Emx1-Cre mice with a conditional halorhodopsin responder line, a technique we previously used to silence S1^27^. We recorded from POm cells while silencing S1 or M1 with an amber laser (Figure 3a). Here, we were similarly able to inhibit M1 activity (Figure 3b).

**Figure 3.**
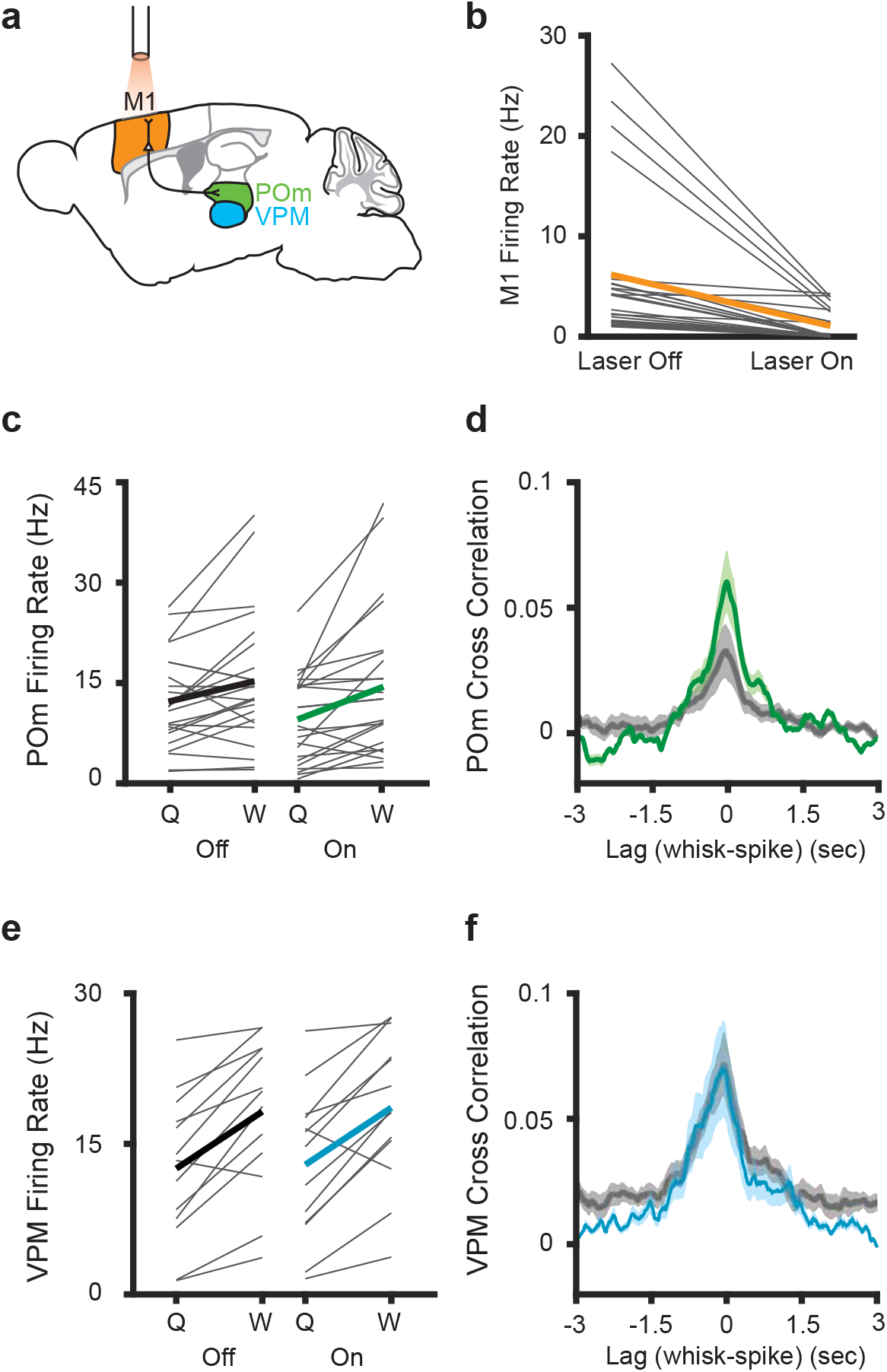
Inhibition of primary motor cortex increases POm correlation with whisking. **(a)** Experimental setup. M1 was optogenetically silenced while recordings were made from M1, POm, or VPM. Adapted from *The Mouse Brain Atlas in Stereotaxic Coordinates*^57^. **(b)** Effect of laser on M1 activity (n = 26 M1 cells, 2 animals, mean decrease of 5.1 Hz, or 83%, p =0.0005). **(c)** Individual (gray) and mean (black or green) POm firing rates during whisking and quiescence when the laser is off or on (n = 23 cells, 3 mice, whisking p = 0.005, laser p = 0.016, two-way repeated measures ANOVA). **(d)** Cross-correlation of POm firing rate and whisking amplitude when the laser is off. The peak correlation was significantly higher when the laser was on (p=0.0018, paired t-test between peak values). **(e)** Individual (gray) and mean (black or blue) VPM firing rates during whisking and quiescence when the laser is off or on (n = 13 cells, 2 mice, whisking p = 0.0002, laser p = 0.27, two-way repeated measures ANOVA). **(f)** Cross-correlation of VPM firing rate and whisking amplitude (p = 0.11, paired t-test between peak values).

M1 suppression reduced the baseline firing rate of POm cells, but POm activity was still elevated during whisking regardless of whether the laser was on or off (Figure 3c). Suppressing M1 increased the correlation between POm firing rate and whisking amplitude (Figure 3d). This suggests that POm encoding of fine whisking kinematics arises from ascending sensory reafference rather than input from motor cortex.

To confirm that these effects were due to inhibition of M1 inputs to POm and not an artifact of optogenetic-induced changes in whisking behavior, we also recorded from cells in VPM. VPM, which does not receive direct projections from M1, was largely unaffected by M1 inhibition. We observed no effect of inhibition on VPM firing rates or cross-correlation between VPM activity and whisking (Figure 3e, f).

In a parallel set of experiments, we silenced S1 using the same cre-dependent halorhodopsin line (Figure 4a). We recently demonstrated that this technique robustly blocks action potentials throughout all cortical layers of S1 in awake behaving mice^27^. Silencing S1 reduced POm activity whether mice were whisking or quiescent (Figure 4b), n = 11 cells, 3 mice; whisking p = 0.0002, laser p = 0.024, two-way repeated measures ANOVA). As in the M1 experiments, the correlation between whisking amplitude and POm activity was, if anything, unchanged or increased by S1 silencing (Figure 4c, laser-off peak correlation = 0.036, laser-on peak correlation = 0.081. There was a tendency for S1 inhibition to reduce overall activity in VPM, possibly reflecting the known corticothalamic connections between S1 and VPM^28–30^, but this effect did not reach significance (Figure 4d; p = 0.1). Suppressing S1 had little impact on the correlation of VPM spiking and whisking (Figure 4e), though there was a trend for S1 inhibition to increase modulation depth (increase from 0.17 to 0.32, 91% change, p = 0.057).

**Figure 4.**
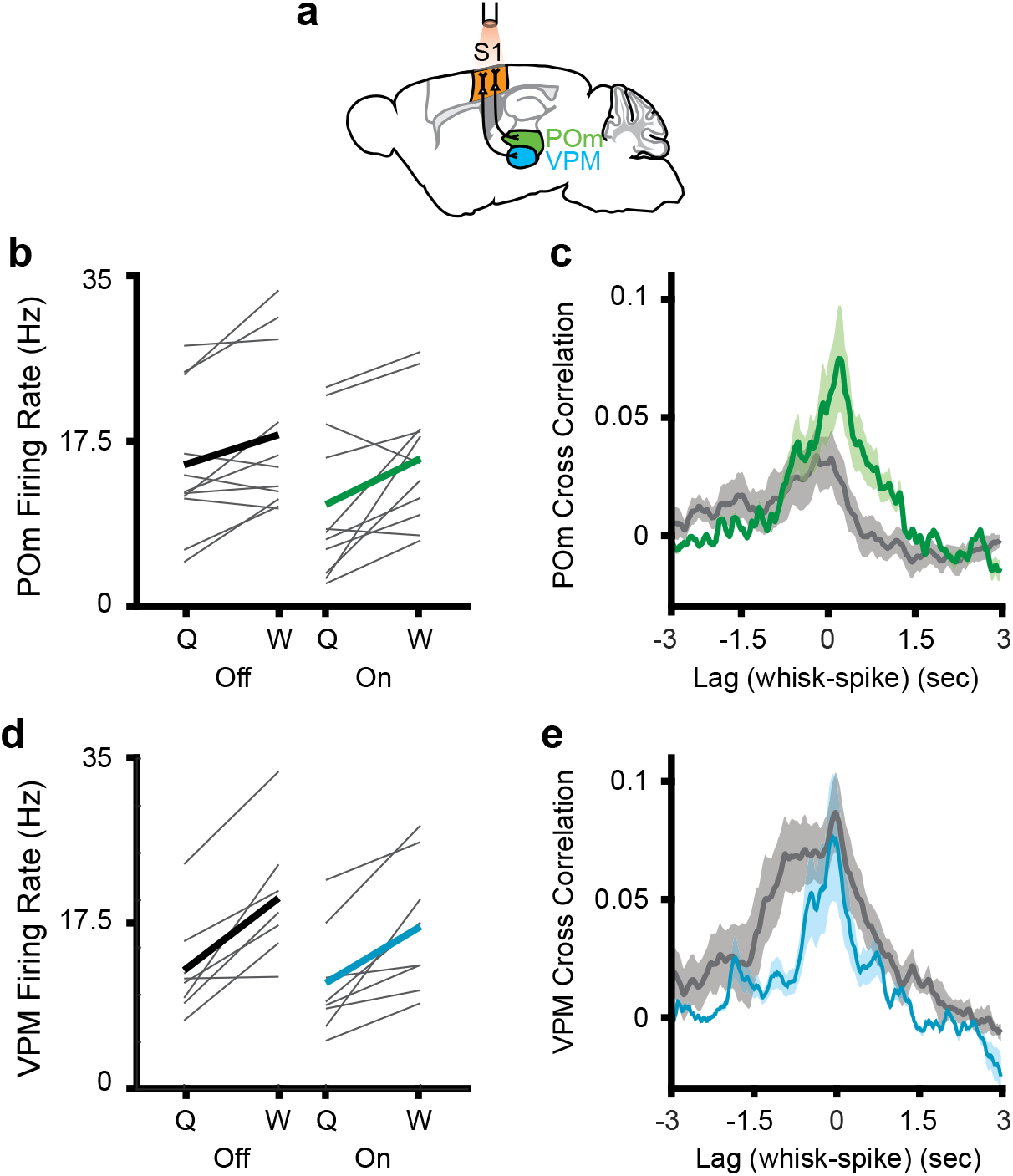
Inhibition of primary somatosensory cortex increases POm correlation with whisking. **(a)** Experimental setup. **(b)** Individual (grey) and mean (black or green) POm firing rates during whisking and quiescence when the laser is off or on. n = 11 cells, 3 mice, whisking p = 0.0005, laser p = 0.03, two-way repeated measures ANOVA. **(c)** Cross-correlation between POm firing rate and whisking amplitude when the laser is off (grey) or on (green). **(d)** Mean VPM firing rate (n = 8 cells, 2 mice. Whisking p = 0.001, laser p = 0.11, two-way repeated measures ANOVA). **(e)** Cross-correlation between POm firing rate and whisking amplitude (p = 0.057, paired t-test between peak values).

Thus, both optogenetic manipulations had qualitatively different effects on VPM and POm activity, consistent with the known anatomical differences in corticothalamic projections onto these two nuclei. Together, these results demonstrate that POm does not inherit information about whisking amplitude from M1 or S1. Rather, corticothalamic inputs appear to transmit signals other than whisker movements, and these additional signals reduce the correlation of POm activity with whisking amplitude and phase.

In addition to afferent inputs from brainstem and efferent inputs from cortex, POm receives excitatory projections from the superior colliculus (SC)^31^, which could also provide a motor efference copy signal similar to known collicular circuits in the visual system^32^. SC receives excitatory input from both the trigeminal brainstem^33^ as well as cortex, making SC a potential source of whisking-related POm activity. To test this possibility, we performed bilateral electrolytic lesions in SC and subsequently recorded POm cells (Figure 5a). Whisking had similar effects on POm activity in both intact and lesioned animals (Figure 5b, n= 49 cells from 8 animals, 59% increase in mean firing rate, p<10^−9^). POm firing rates of lesioned mice were overall higher than those of intact animals, independent of whether animals were whisking or quiescent (Figure 5c, lesion p < 10^−3^, whisking p < 10^−5^, 2-way ANOVA). There was a slight tendency for SC-lesioned animals to whisk more frequently than intact animals, but this effect was not statistically significant (Figure 5d, p = 0.35, Wilcoxon rank-sum test). We conclude that SC is not responsible for the whisking-induced elevation of POm activity.

**Figure 5.**
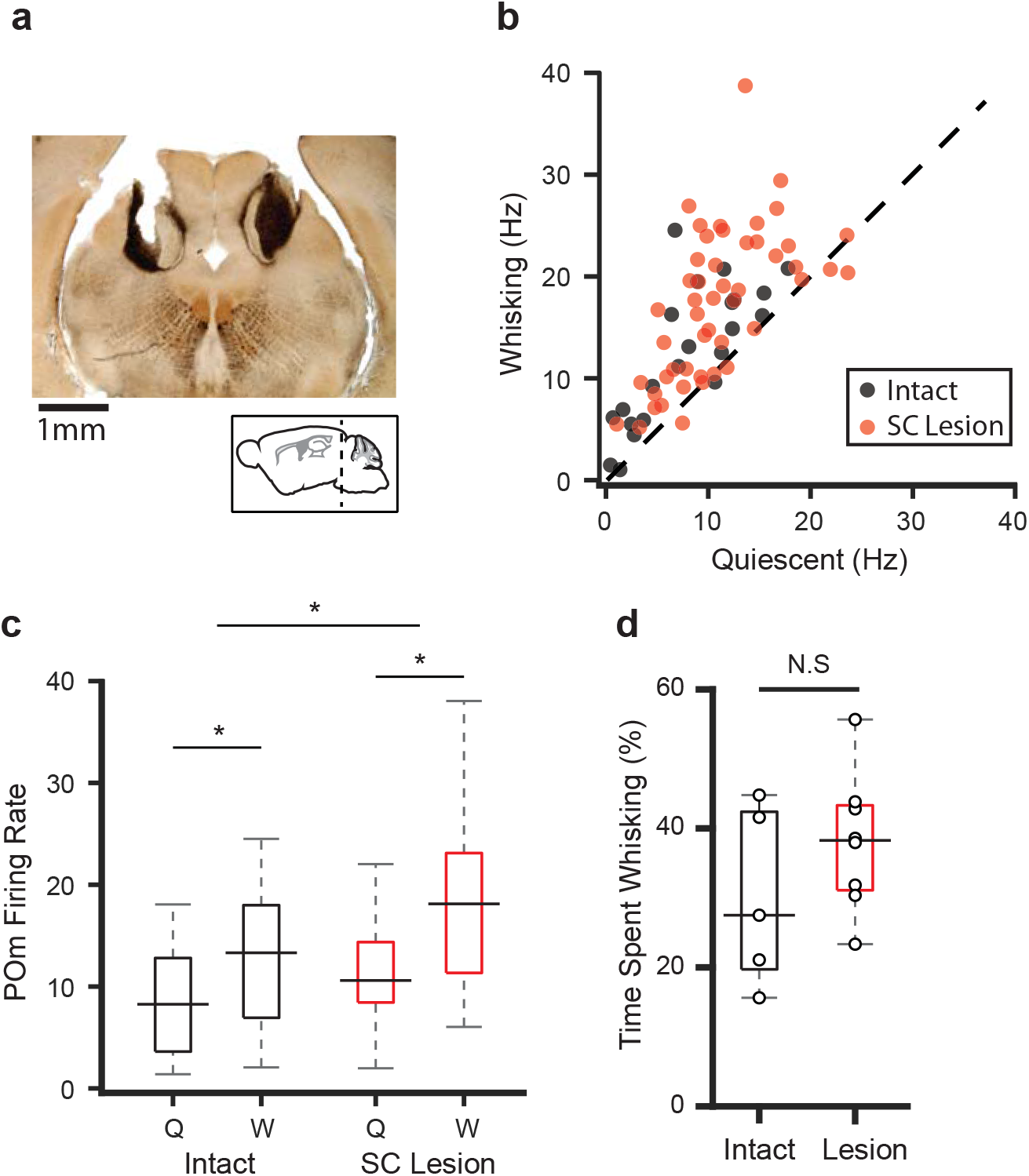
Lesions to superior colliculus do not reduce POm correlation with whisking. **(a)** Sample coronal section showing bilateral electrolytic lesion of superior colliculus. **(b)** Scatter plot of POm firing rates during whisking and quiescence in lesioned (red) and intact animals (black, data from Fig. 1). Firing rates in lesioned animals were significantly higher during whisking (n = 49 cells from 8 animals, increase from mean of 10.9Hz to 17.4Hz, or 59%, p < 10^−9^, paired t-test). **(c)** Box plots of POm firing rates during whisking (W) and quiescence (Q) in intact (black) and lesioned animals (red). Pom firing rates in lesioned animals were higher than intact animals (whisking p < 10^−5^, lesion p < 10^−3^, 2-way ANOVA). **(d)** Lesioned animals tended to spend slightly more time whisking, but this was not statistically significant (intact median = 27.5%, n = 5 mice; lesion median = 38.3%, n = 8 mice, p = 0.35, Wilcoxon rank-sum test).

Neither reafference nor the most likely sources of motor efference copy explain the coarse modulation of POm by movement. This raises the question of whether POm encodes movement *per se,* or another variable that is coupled with whisking and other movements, such as arousal. To investigate this, we measured pupil diameter, which is a known metric of arousal. We acquired videos of the pupil and whiskers while recording from POm (Figure 6a). Pupil diameter was tightly correlated with whisking, with pupil dilation lagging whisking amplitude by 880 msec on average (Figure 6b). Pupil diameter also correlated with POm activity, to a similar degree as whisking and with a lag of 950 msec (Figure 6c, whisking amplitude peak correlation = 0.052, pupil diameter peak correlation = 0.071, p = 0.23, paired t-test).

**Figure 6.**
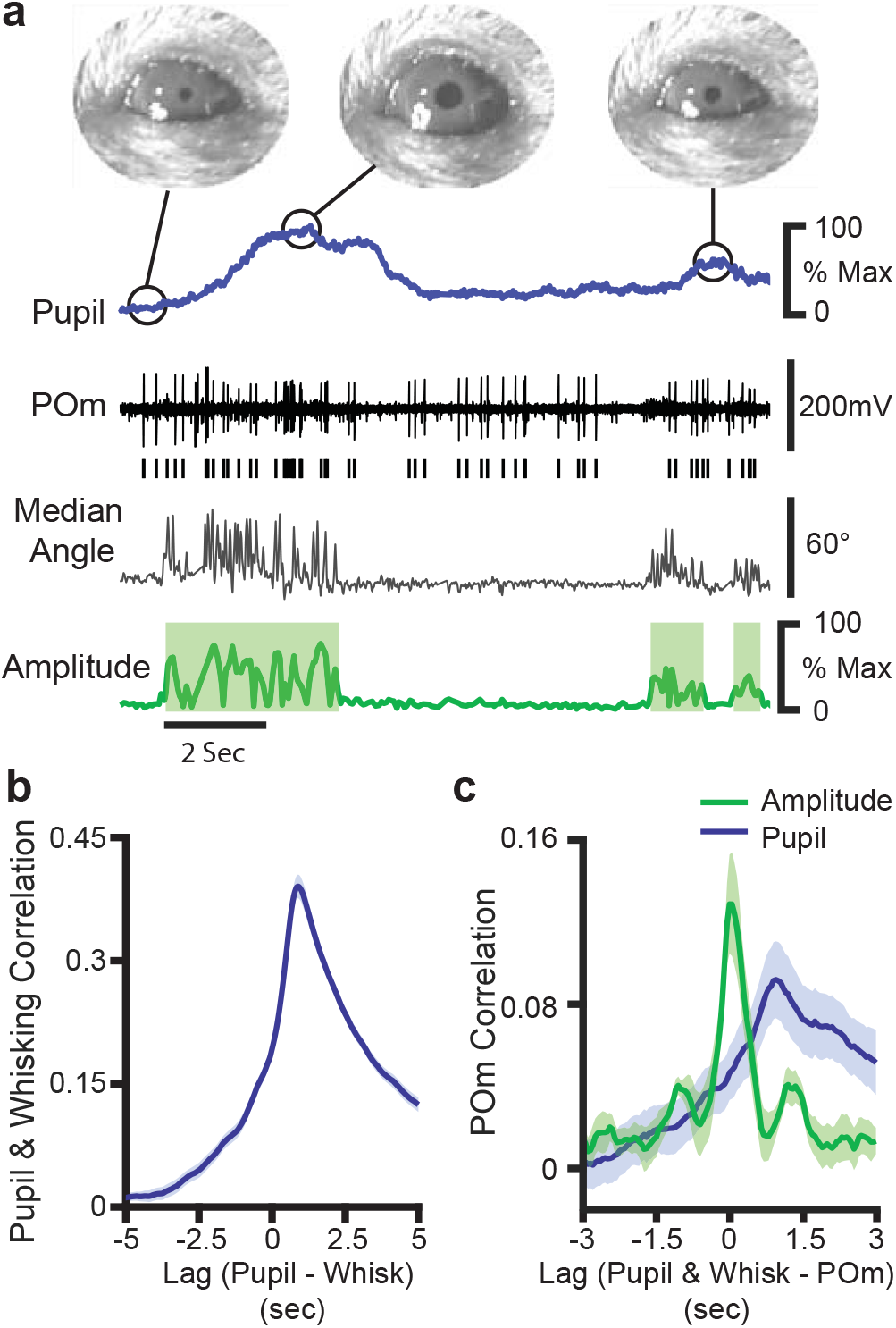
POm activity tracks pupil dynamics. **(a)** Sample recording of POm activity (middle, black) with concurrent ipsilateral pupil diameter (blue, top), median whisker angle (*middle, gray*), and whisking amplitude (green, bottom). **(b)** Cross-correlation of pupil diameter and whisking amplitude (30 recording sessions from 7 animals). Errors bars are present but very small. **(c)** Cross-correlation of POm firing rate (n = 10 cells from 3 animals) with whisking amplitude (*green*) and pupil diameter (*blue*).

We reasoned that, if the modulation of POm by whisking was truly due to whisker movement rather than some other correlated variable, non-somatosensory thalamic nuclei would not be expected to track whisking. The secondary visual thalamic nucleus LP is the rodent homolog of the primate lateral pulvinar. LP is primarily coupled with cortical and subcortical visual areas^34^, rather than somatosensory ones. Because of their different connectivity, POm and LP are expected to carry separate sensorimotor signals related to somatosensation and vision, respectively. Therefore, LP would not be expected to encode whisker movement. By contrast, changes in behavioral state, such as overall animal arousal as suggested by our pupil measurements, might modulate all thalamic nuclei, including LP and POm.

We tested this idea by recording juxtasomally from LP neurons (Figure 7a, b; 29 cells from 4 mice). Surprisingly, we found that LP activity was significantly increased during whisking bouts (Figure 7c, increase from 13.0Hz to 18.0Hz, p < 10^−4^, paired t test). Like POm, LP activity correlated with both whisking amplitude and median whisker angle with low latency (Figure 7d). Since changes in pupil diameter will cause more light to fall on the retina, the LP correlation with whisking might be an artifact of pupil dilation. To control for this, a subset of cells were recorded in low light. Under these darker conditions, the pupil was maximally dilated and did not change (Figure 7b), rendering input to the retina largely constant. However, these cells still showed an equivalent increase in firing rate during whisking (Figure 7c, orange; n = 29 cells, 4 animals; increase from a mean of 13.0Hz to 18.0 Hz, or 39%, p < 10^−4^, paired t-test). Thus, LP activity appears to track whisking independent of changes in visual input, which suggests that the effect in both nuclei is due to the arousal-whisking correlation rather than a direct effect of whisking.

**Figure 7.**
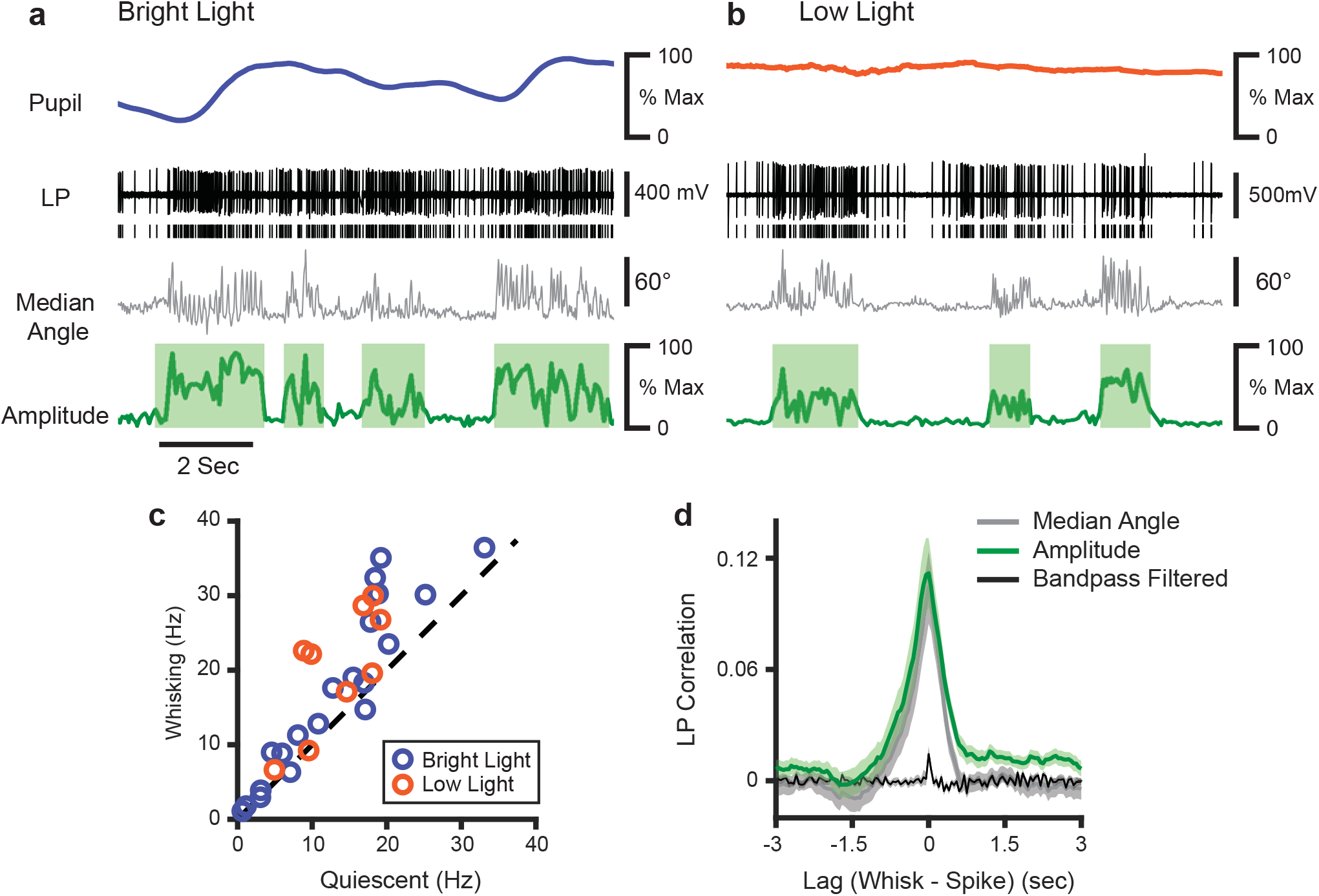
LP activity tracks slow whisker dynamics. **(a,b)** Sample recordings of two LP cells (black) recorded in normal light **(a)** or low light **(b)**, with corresponding median whisker angle (*gray*) whisking amplitude (green), and pupil diameter (blue or orange). **(c)** Scatter plot of mean firing rate in LP cells during whisking and quiescence. *Blue*, cells recorded in bright light; *Orange*, cells recorded in low light (p < 10^−4^, paired t-test). **(d)** Cross-correlation of LP firing rate with whisking amplitude (green), median whisker angle (red), and 4−30 Hz bandpass filtered angle (black).

Together, our results indicate that the slow component of whisking-related activity in POm is neither a consequence of ascending motion signals from reafferent mechanisms nor corticothalamic or colliculothalamic efferent mechanisms. We conclude instead that behavioral state, such as arousal, may strongly dictate the activity of secondary thalamic nuclei, including POm and LP.

## Discussion

Our study tested the idea that secondary somatosensory thalamus is a monitor of movements or motor commands and manipulated the multiple known pathways to POm that could mediate such signals. Juxtasomal recordings of POm cells revealed that this nucleus mainly tracks slow components of whisking, not detailed kinematics. Consistent with other studies^8,17^, mouse POm firing rates are much higher during bouts of whisking than when a mouse is quiescent. However, POm activity mainly correlates with the slow change in whisking amplitude rather than the fast changes of the whisk cycle. Only a minority of our POm cells exhibited any whisking phase information, and phase encoding appeared to depend on sensory reafference. We have demonstrated that, by contrast, the overall elevation of POm activity by whisking is not due to sensory reafference from self-generated movements, as transection of the facial motor nerve did not uncouple POm activity from ipsilateral whisking. We showed that potential motor efference copy via corticothalamic pathways from S1 and M1 cannot account for whisking modulation of POm. Similarly, the phenomenon is independent of superior colliculus, the activity of which is linked to movement and orienting.

What appears to be movement-related activity in POm is likely instead a consequence of the encoding of behavioral state. Activity in secondary visual thalamus (LP) exhibits the same correlation with whisker movement that we observed in POm. Though it is possible that POm and LP separately encode correlated sensorimotor information, a more parsimonious explanation is that both POm and LP are modulated by arousal, which is naturally elevated during movement. Modulation of activity by behavioral state may be a general property of all secondary thalamic nuclei. Future studies are needed to examine if this principle holds in auditory thalamic subnuclei and perhaps even thalamic nuclei connected to motor cortex and other frontal areas. Conceivably, some movement correlations seen even in motor thalamus^35^ may reflect various states more than specific movements.

The paralemniscal system has been speculated to be a parallel secondary afferent pathway^16,19^. However, in anesthetized rats, POm does not appear to be sensitive to fine aspects of whisker touch, having very large receptive fields and long-latency responses^9,13^. One might expect that very large synchronized movements of the whiskers, such as during whisking, would elicit a response from POm due to sensory reafference driving coarse receptive fields. However, paralyzing the face did not uncouple POm activity from ipsilateral whisking amplitude (Figure 2). Similarly, mouse barrel cortex is also modulated by whisking and quiescence in absence of sensory input: whisking is associated with a decrease in synchrony between layer 2/3 pyramidal cells in S1 and an increase in discharges by VPM, which is unaltered by bilateral transection of the infraorbital nerve sensory nerve^36^. Manipulations of somatosensory thalamus strongly impacted cortical synchronization^37^. Further studies are needed to parcel out the extent to which thalamic contributions to cortical synchronization is due to inputs from VPM, POm, or both.

POm receives descending input from many cortical regions including M1 and S1. Conceivably these inputs could modulate ascending sensory input or provide the thalamus with a motor efference copy^15,38^. Similarly, LP and LGN axons in V1 exhibit eye movement-related signals^23^. Previous studies in anesthetized rats have shown that cortical inactivation will silence POm, but not VPM^9^. Therefore, cortex might be the primary source of excitatory input to POm. However, we discovered that, in the awake mouse, silencing either M1 or S1 only slightly reduces the firing rate of POm cells and has little to no effect on VPM activity (Figure 3). We conclude that, while S1 and M1 provide significant excitatory inputs to POm, these inputs are not the sole drivers of POm activity during wakefulness.

Moreover, silencing these corticothalamic pathways increased rather than decreased the correlation between POm activity and whisking amplitude. If POm activity were primarily representative of a cortical efference copy, we would expect the opposite effect. While we cannot rule out the possibility that POm receives some efference copy from cortex, such input is not the cause of what at first appears to be whisking modulation. POm might instead be under equal or greater control of subcortical regions such as trigeminal brainstem complex, zona incerta, the thalamic reticular nucleus, and neuromodulatory brainstem centers – all of which receive inputs from broad areas of the nervous system^13,39^.

As POm continues to track whisking in absence of both ascending sensory input and descending cortical input, we propose that the activity we observe is not sensorimotor in nature, but rather representative of thalamic coding of internal state. POm axons project to the apical dendrites of pyramidal cells^6,40^, where they might drive state-dependent changes in activity and synchrony. Arousal has dramatic effects on cortical dynamics^41–43^. We observed that pupil diameter, which closely tracks arousal, is highly correlated with whisking amplitude. Due to the coupling between pupil and whisking dynamics, they both correlate with POm firing rates (Figure 6). To dissociate the contributions of arousal and whisker movement, we took the novel approach of comparing POm dynamics with those of LP, the rodent homolog of the primate lateral pulvinar. We found a near-identical relationship between LP activity and whisking as we observed in POm (Figure 7), even though there is no known connectivity between LP and the whisker system. As for POm, these shifts in LP activity do not appear to be sensory dependent, as they persist even in low-light conditions where the pupil is maximally dilated and can no longer contribute to changes in retinal activity.

If state-dependent modulation of secondary thalamic nuclei is not derived from sensory reafference or motor efference copy from cortex or superior colliculus, the likely remaining candidates would include a large number of neuromodulators. For instance, zona incerta terminals within POm are regulated by acetylcholine^44^ and are likely modulated in the same way within LP. However, acetylcholine and norepinephrine both track pupil dynamics^45^, and both are also plausible mechanisms. In addition to these two well-studied modulators, there are many others known to have functions in thalamus^46^. Furthermore, any of these modulators could act directly on POm and LP or indirectly through ZI, TRN, brainstem nuclei, or other inputs.

The arousal effect we have described may be a more general version of modality-specific attentional effects that have been proposed for at least some secondary thalamic nuclei. In primates, pulvinar neurons respond strongest when stimuli are presented in attended regions of visual space^47^, and lesion of the pulvinar leads to deficits of selective attention during visual tasks^22,48^. Human patients with pulvinar damage exhibit spatial neglect, in which a stimulus can be perceived normally in isolation but is missed or distorted in the presence of neighboring stimuli^49,50^. By analogy, one might hypothesize that POm provides feedback that selects somatosensory stimuli for further cortical processing. Indeed, we and others have already demonstrated that activation of POm sensitizes cortical pyramidal neurons to the occurrence of subsequent tactile stimuli^14,15^. Thus, POm affords control over the gain of the sensory responsiveness of somatosensory cortex circuitry. Selective enhancement of sensory responses by attention within a modality could be a general principle of all secondary thalamic function.

Cortex-wide fluctuations in activity are known to correlate with various uninstructed movements^51^ Cortical activity ceases in the absence of thalamic input^35,52^, and secondary thalamic inputs to somatosensory cortex are stronger and longer lasting than corticocortical connections^14^. Taking those studies and our study together suggests that secondary thalamus may be the underlying cause of the recently observed patterned fluctuations in activity across cortex. Our study directly tested the multiple known possible sources of afferent and efferent motor signals to secondary thalamus. None of these could explain apparent shifts in thalamic activity. Thus, behavioral state, rather than uninstructed movement, may be a primary driver of thalamic and cortical activity during movement.

Elevated firing rates in secondary thalamus due to arousal or attention could be useful for creating periods of heightened cortical plasticity. Recent studies have shown that repetitive sensory stimuli in anesthetized animals drives POm input to pyramidal neurons, which leads to enhancement of future sensory responses in cortex^53^. A potential mechanism of this is that disinhibition of apical dendritic spikes leads to long-term potentiation of local recurrent synapses among cortical pyramidal neurons^54^. Furthermore, an *in vivo* study found that associative learning can also potentiate long-range POm connections onto pyramidal neurons when subsequently measured *in vitro*^55^.

It is conceivable that the arousal modulation of secondary thalamus that we have described is utilized by such processes. Our work opens avenues to examining potential links between arousal, attention, and plasticity.

## Acknowledgments

The authors thank C. Kellendonk, A. Das, C. Rodgers, S. Benezra and G. Pierce for comments on the manuscript and D. Baughman, B.C. Pil, and L. Jordan for technical support. This work was supported by NIH R01 NS094659, R01 NS069679, and T32 EY013933.

## Author Contributions

Conceptualization: AKK and RMB; Investigation: AKK and GHP; Formal Analysis: AKK, GHP, and RMB; Optogenetics Methodology, Resources, and Assistance: YKH; Writing: GHP and RMB.

## Methods

All experiments complied with the NIH Guide for the Care and Use of Laboratory Animals and were approved by the Institutional Animal Care and Use Committee of Columbia University. Twenty-two C57BL/6 mice were used in these experiments.

### Surgery

Mice were anesthetized with isoflurane and placed in a stereotax. The skull was exposed, a thin layer of superglue was applied, and a custom-cut stainless steel headplate was attached using dental acrylic. A small (200 μm wide) opening was made on the mouse’s left side at ~1.7 mm posterior to bregma and 1.4 mm lateral of the midline. A silver wire or screw was inserted over the frontal cortex of the same hemisphere as a ground electrode and covered with dental acrylic. The skin was sealed to the implant using superglue. Mice were allowed to recover from surgery for 5 days before habituation. Mice were habituated to the setup for 5 days by attaching their headplate to a holder on the recording table for 5−30 min each day, during which no recordings were performed.

### Electrophysiology

After habituation, a mouse would be recorded from for 3−7 days. A glass micropipette (opening ~1.5 μm ID, shank ~60−80 μm OD over last 3−4 mm) was filled with artificial cerebrospinal fluid (aCSF) and inserted vertically into the brain using a micromanipulator. POm cells were typically recorded at microdrive depths of 2800−3600 μm relative to the pia, and LP cells were recorded at depths of 2100−2600 μm relative to pia. Recordings were made with a MultiClamp 700B amplifier (Molecular Devices), bessel filtered 300−10,000 Hz, and digitized at 16 kHz using custom Labview software (ntrode). At the end of some experiments, recording sites were labelled with a glass electrode coated in DiI inserted to a depth of 3600 μm relative to the pia.

### Videography

Whisker and pupil videos were made during electrophysiology and imaging using multiple PS3eye cameras running at 125 frames per second. Camera housings had been removed, and the lenses replaced with a 12 mm F2.0 lens (M12 Lenses Inc, part # PT-1220). Video was acquired using the CodeLaboratories PS3eye camera driver and the GUVCView software on linux computer.

### Optogenetics

Optogenetic silencing of cortex was performed using Emx1-Halo mice as previously described ^27^. Briefly, Emx1-IRES-Cre knock-in mice (Jackson Laboratories, stock #005628) were crossed to Rosa-lox-stop-lox (RSL)-eNpHR3.0/eYFP mice (Ai39, JAX, stock# 006364), which express halorhodopsin after excision of a stop cassette by Cre recombinase. All mouse lines were maintained on a C57BL/6 background. Optogenetic experiments used mice that were heterozygous for the desired transgene as assessed by in-house genotyping. The locations of S1 and M1 were marked based on stereotaxic coordinates during headplate surgery, and the skull was thinned before recordings. Light was generated by a 593- or 594-nm laser (OEM or Coherent) coupled to a 200-μm diameter, 0.39 NA optic fiber (Thorlabs) via a fiberport, and the diamond-knife cut fiber tip was placed above M1 or S1.

### Nerve Transection

The facial nerve was transected with the mouse under isoflurane anesthesia. A small (~5 mm) incision, centered ~5−8 mm ventral of the eye, was made in the skin. The buccal branch of the facial was identified running from near the ear to the whisker pad, blunt dissected free of underlying tissue, and cut. The skin was closed with stitches and bupivacaine applied.

### Superior Colliculus Lesion

The superior colliculus was lesioned bilaterally just prior to headplate implantation. Craniotomies were drilled over superior colliculus (0.5 mm anterior of lambda, 0.75 mm lateral of midline). A tungsten electrode (0.3−1.0 MΩ) was inserted to depths of 1 mm and 2 mm on each side, and 300 μA of current was delivered for 30 s at each lesion site. Mice were then implanted with a headplate and habituated as described above. Histology was used to confirm lesion size and location, and only recordings from mice with on-target lesions were analyzed.

### Histology

At the end of experiments, mice were deeply anesthetized with sodium pentobarbital and then perfused transcardially with 1× phosphate buffer followed by 4% paraformaldehyde. Brains were removed and sectioned on a vibratome into 100 μm-thick slices, or on a freezing microtome into 50 μm-thick slices. 100-μm slices were mounted directly on glass slides with mounting medium. 50-μm slices were stained in a solution of Cytochrome C (0.3 mg/ml), Catalase (0.4mg/ml), and 3−3’-Diaminobenzidine (DAB, 0.583mg/mL). Sections were incubated in this solution at 40°C for 30−45 minutes. Sections were washed 5 times in 1× phosphate buffer and mounted on glass slides with mounting medium.

### Data Analysis

Putative action potentials were identified offline with custom MATLAB software. Spikes were then manually sorted with MClust (version 4.3).

Whiskers were automatically tracked from videos using software (Clack *et al.* 2012). Custom MATLAB software was used to compute the median whisker angle. The median angle was bandpass filtered from 4 to 30 Hz and passed through a Hilbert transform to calculate phase. We defined the upper and lower envelopes of the unfiltered median whisking angle as the points in the whisk cycle where phase equaled 0 (most protracted) or π (most retracted), respectively. Whisking amplitude was defined as the difference between these two envelopes. Periods of whisking and quiescence were defined as times where whisking amplitude exceeded 20% of maximum for at least 250 msec. Periods of time where amplitude exceeded this threshold for less than 250 msec were considered ambiguous and excluded from analysis of whisking versus quiescence.

For cross-correlation analysis, whisking angle, amplitude, pupil, and spike vectors were binned with a 10-millisecond time bin. They were then normalized to have a mean of zero and standard deviation of one. Cross-correlations were again normalized such that the autocorrelation at a time lag of zero equaled one. To test the significance in changes between cross-correlation distributions (*e.g.* when comparing laser-off and laser-on conditions during cortical silencing) we found the lag of the peak correlation value for each distribution. We then performed paired t-tests between the correlation values of each cell at that time lag.

For each cell, each spike that occurred while the mouse was whisking was assigned a phase. The distribution of possible spike phases (-π to π) was calculated using 32 equally sized bins. Using the same binning, we then calculated the distribution of phases observed in the video to determine the time the whiskers spent at various mean phases. We then normalized the spike phase distribution by the phase distribution to calculate firing rate as a function of phase. The modulation of the cell was characterized by fitting a sine function with a period of 2π to this rate function using least-squares regression. The modulation depth was calculated as the amplitude of the fitted sine wave divided by the cell’s mean firing rate as in8 ^8^. To test the significance of this modulation, we compared the distributions of whisking phase and (unnormalized, unbinned) spike phase with a Kuiper test and a Bonferroni multiple-comparisons correction.

Pupil diameter was measured from video using custom MATLAB software. Videos were level-adjusted and thresholded to maximize the contrast between the pupil and the rest of the eye. The built in imfindcircles() function was used to locate the pupil and measure diameter on each frame.

